# Structural Analysis of Recombinant AAV Vector Genomes at Single-Molecule Resolution

**DOI:** 10.64898/2025.12.04.692441

**Authors:** David Rouleau, Dimpal Lata, Serena Dollive, Robert E. Bruccoleri, Laura Van Lieshout, Diane Golebiowski, Ify Iwuchukwu

## Abstract

Recombinant adeno-associated virus vectors are essential tools for *in vivo* gene therapy, yet heterogeneity in their packaged genomes remains an important safety consideration. To systematically evaluate this heterogeneity, we developed a long-read, read-level analysis pipeline that directly classifies individual AAV genomes and their structural variants from PacBio sequencing data.

The workflow combines two components: a tiling step that aligns each read to reference sequences to generate positional patterns, and a parsing step that applies a formal grammar to categorize reads into five structural classes: expected, truncated, snapback, truncated snapback, and others. Each molecule is annotated with strand orientation, breakpoint coordinates, and structural arrangement, enabling precise classification of genome heterogeneity at single-vector resolution.

Applied to both single-stranded and self-complementary vector genome preparations, the pipeline achieved high classification accuracy and revealed distinct patterns of genome structure between different vector constructs. In both cases, the majority of genomes were classified as expected full-length species, consistent with the dominant full peaks observed by orthogonal methods. For snapback genomes, breakpoints frequently clustered at discrete sites, with some coinciding with regions predicted to form stable secondary structures and others occurring in less structured regions. This distribution suggests contributions from both sequence-driven folding and additional replication- or processing-related mechanisms. Together, these read-level insights highlight sequence and structural features that shape AAV genome heterogeneity. Importantly, the pipeline demonstrated strong performance in structural classification, maintaining high accuracy even in the presence of sequencing error profiles such as homopolymer-associated indels (insertion or deletion).

By integrating structural classification, sequence context, and secondary-structure predictions, our pipeline provides a comprehensive framework for evaluating recombinant adeno-associated virus genome diversity. This approach not only improves resolution of vector genome architecture but also offers actionable insights to guide vector design and production processes for safer and more efficacious recombinant adeno-associated virus therapeutics.

## 1. Introduction

Recombinant adeno-associated vectors (rAAVs) are emerging as a valuable tool for their use in gene therapy, with several candidates showing promising clinical trial outcomes and some already approved by the U.S. Food and Drug Administration (1–3). At the same time, clinical experience has highlighted important safety concerns including hepatotoxicity, thrombotic microangiopathy, myocarditis, and hepatocellular carcinoma (4–7). In some high-dose systemic trials, such as those for X-linked myotubular myopathy, patient deaths have been linked to acute liver failure and immune-related complications (8). More recently, the Elevidys trial for Duchenne muscular dystrophy was temporarily halted after multiple patient deaths, drawing renewed attention to dosing and safety thresholds (9). Mechanistic studies suggest these outcomes may be driven by innate and adaptive immune responses to AAV capsids and transgene products, off-target biodistribution of vector genomes, and, in rare cases, insertional mutagenesis leading to oncogenic risk (8,10). These observations emphasize the importance of maintaining the highest possible quality of rAAV vector genomes, as improved potency at lower doses may help reduce immune responses and off-target effects

Achieving high-quality rAAV preparations, however, remains challenging due to the frequent presence of noncanonical vector genomes, which have been linked to unintended biological effects in vitro and *in vivo* (11). Another obstacle is the presence of canonical genome structural variation including snapback AAV (12–14), truncated AAV (15,16) and self-priming truncated species (17). These variants arise during AAV replication and packaging and have been observed both in production samples and *in vivo* (18–20). These structural variants, despite only containing DNA of the desired vector genome product, can also have a negative impact on vector productivity (21,22). Thoroughly characterizing the full spectrum of vector genome species in rAAV preparations is therefore essential for both safety and efficacy.

Short read sequencing methods are unable to robustly resolve all structural genome rearrangements present in AAV samples such as truncation products due to the need for *in silico* read assembly (12,19,23). Recent advances in long-read sequencing technologies, such as single-molecule real-time (SMRT) sequencing, have enabled the direct capture of complete AAV vector genomes without the need for read assembly (24). Using single-molecule, real-time next-generation sequencing (SMRT NGS) via the PacBio Sequel II, we captured complete AAV vector genomes via long read sequencing without the need to assemble any reads bioinformatically. This approach gave us a snapshot of the distribution of structural variants present within the recombinant virus population. Since AAV vector characterization is a relatively unique use case, publicly available software that can interpret these full AAV sequences and report structural variant populations are sparse (25,26). Here we introduce a comprehensive analysis pipeline designed to evaluate the molecular structure of recombinant vector genomes at single-vector resolution.

Our pipeline is made up of two programs: tiling and subparsing. Tiling, described previously (27), uses NCBI-BLAST (28) to break sequence reads into a series of alignments that best cover as much of a read as possible. AAV genomes with noncanonical DNA, such as helper plasmid DNA or gene-of-interest plasmid backbone DNA, can be counted in the tiling file as the sequences with BLAST alignments to those sequences. To categorize sequences with only canonical AAV genome alignments, we use the second part of our pipeline, the subparser, which is the focus of this paper.

Our subparser program assigns canonical AAV genome sequences to one of sixteen structural variant categories (Fig. 1) which can be grouped into five broader morphological classes: expected, truncated, snapback, truncated snapback, and other genome forms. Several of these categories are labeled as self-priming variants. Self-priming refers to AAV genome molecules in which the 3′ inverted terminal repeat folds back to form a double-stranded hairpin that primes complementary-strand DNA synthesis (29). We use the term self-priming rather than self-complementary because such structures can also arise during PacBio SMRTbell library preparation, where single-stranded AAV vector may undergo partial strand fill-in reactions that generate molecules resembling a self-complementary vector (30).

**Fig 1.**
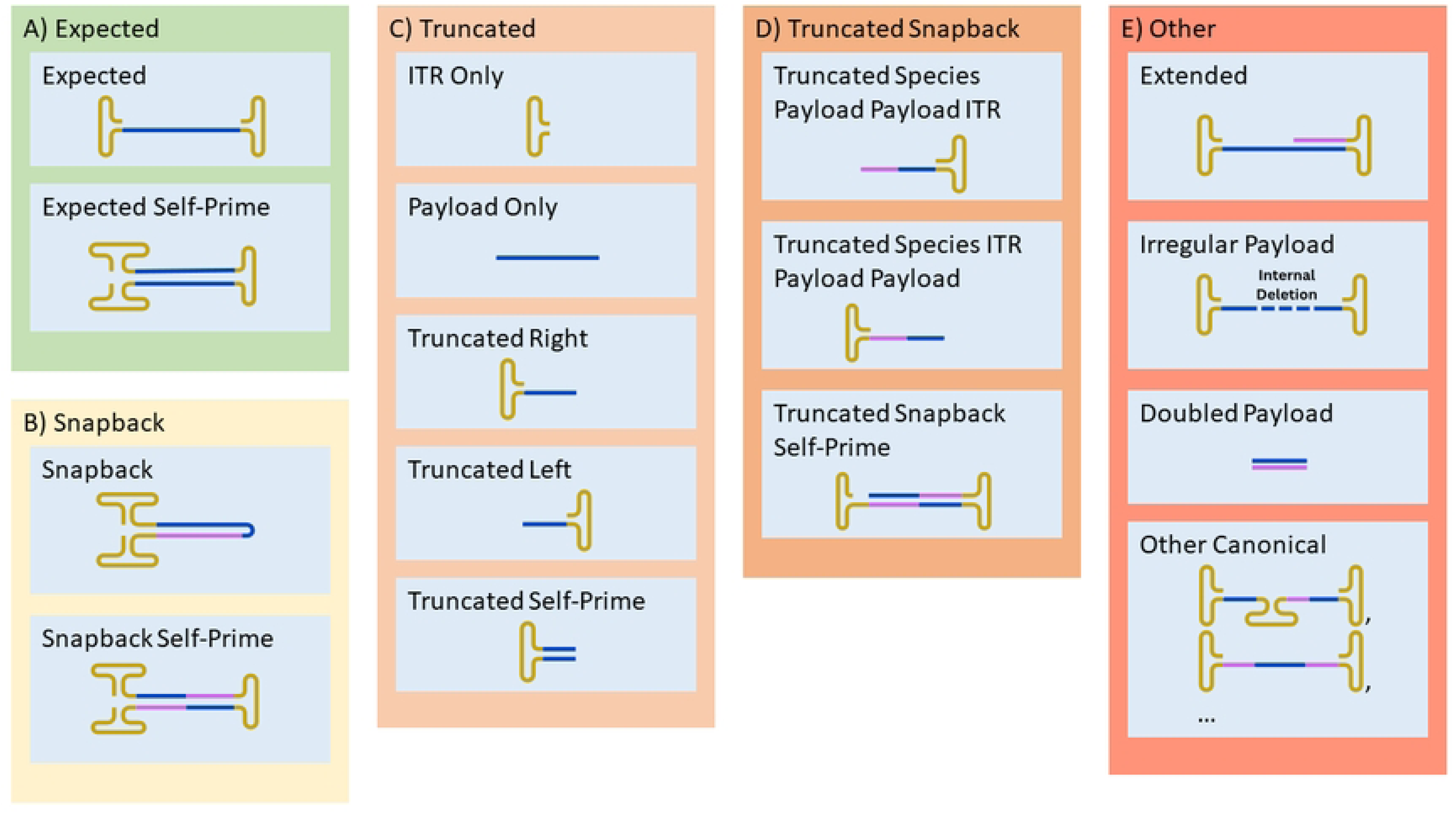
AAV genome structural variants categorized by our subparser program: expected, truncated, snapback, truncated snapback, and other Gold regions of the vector genome represent ITR sequences, and blue and pink regions represent adjacent non-continuous payload (the AAV’s transgene or gene-of-interest region located between the two ITRs) sequences. A) Expected vector genomes represent the full-length, intact recombinant vector genome, spanning from one inverted terminal repeat (ITR) to the other, and containing the complete payload as designed. B) Snapback genomes are formed when single-stranded AAV DNA folds back on itself via internal complementarity, resulting in partially double-stranded molecules with hairpin-like structures. C) Truncated genomes are incomplete vector genomes that have lost portions of their sequence, typically due to replication or packaging defects (31). D) Truncated Snapback Genomes are a snapback genome which have also lost portions of their sequence. E) Other vector genomes include miscellaneous structural variants, including extension products resulting from unresolved ITR termination during genome replication (17,32,33). Four structural variants are labelled as self-priming variants. This different term is used instead of self-complementary for ambiguity since these kinds of structural variants can be present in ssAAV samples due to the DNA repair step of the PacBio SMRT NGS library prep process (Fig A in S1 File)

The subparser is designed to output sequences with different structural variant classifications into different files so that sequences can be filtered based on their category for further downstream analysis. To further study snapback and truncated structures, which are observed to have variance in AAV preparations (15,32), we developed a custom analysis pipeline to analyze snapback and truncated AAV genome sequence files output by the subparser, identify strand-specific breakpoints in vector genomes, and examine local sequence structure. The full implementation and extension of this method are also described in this paper.

## 2. Materials and Methods

### 2.1 AAV vector sample

The vectors analyzed in this study were recombinant GFP constructs, including a self-complementary GFP (scGFP) containing a genome of approximately 2080 nucleotides (nt) with a payload size of 1831 nt, and a single-stranded GFP (ssGFP) containing a genome of approximately 2162 nt with a payload size of 1872 nt. Both vectors were manufactured at Oxford Biomedica (Bedford, MA, USA) using triple-plasmid transfection in HEK293 cells and purified using standard column-based methods.

### 2.2 NGS Library Prep

To assess encapsidated vector genomes (scGFP and ssGFP), DNA was extracted from enriched AAV fractions using the PureLink™ Viral RNA/DNA Mini Kit (Invitrogen) with an added DNase treatment to remove unencapsidated DNA. DNA quality was evaluated using TapeStation (Agilent), Nanodrop, and Qubit (Thermo Fisher Scientific). SMRTbell® libraries were then prepared according to the PacBio DNA amplicon library preparation protocol (PacBio, Menlo Park, CA), which involved ligation of hairpin adapters to generate circular templates suitable for sequencing. Libraries were purified, quantified, and loaded on the PacBio Sequel II system. Data were initially processed using PacBioSMRT Link v11.1 analysis software, and bi-stranded CCS (circular consensus sequence) calling was performed to generate high-accuracy reads for downstream structural variant analysis. Sequencing results were analyzed with our NGS structural variant analysis pipeline. Sequencing results were analyzed with our NGS structural variant analysis pipeline.

### 2.3 AUC Analysis

To assess the accuracy of the results of the subparser on real vector genome samples (scGFP and ssGFP), we ran two vector samples through our NGS structural variant analysis pipeline and through Analytical ultracentrifugation (AUC) analysis and compared the results (15,34). AUC was performed using an Optima AUC instrument (Beckman Coulter) to determine the proportion of full, intermediate, and empty capsids in each AAV sample. Sedimentation profiles were analyzed with the SEDFIT c(s) model, which generates distributions of sedimentation coefficients corresponding to different capsid populations. Integration of the peaks provided the relative percentage of each species within the samples.

### 2.4 Structural Variant Calling Software Pipeline

#### 2.4.1 Tiling

The tiling algorithm, described in detail previously (27), is summarized here for methodological completeness. The tiling algorithm derives its name from its approach of using BLAST to break full sequence reads down into a list of local alignments, the results of which are formatted into space-delimited strings of text called tiles. Each “tile” gives a single BLAST result with the name of the subject sequence that the read segment aligned to, the coordinates of that subject sequence that was aligned to, and the orientation of the alignment. These local alignments are calculated with the read as the query sequence and a list of reference sequences as the subject sequence database. The reference sequences that can be used include the transfection plasmid used to generate the AAV vector product, the host cell genome, and any number of helper plasmids used in vector production.

After many local alignments are generated for a sequence, the tiling algorithm picks up the ordered subset of these alignments that best cover the entire sequence read. The result of this is called a tile pattern, and the occurrence count of each tile pattern within a sample is calculated. The final output of the tiling algorithm, a space separated formatted file labelled as a *.tile.counts file, contains three columns of data for each tile pattern in the sample: the occurrence count, the proportion of reads in the sample that matched the tile pattern, and the tile pattern itself (Table 1). In reasonably homogenous samples with more than 100,000 sequence reads, thousands to tens of thousands of tile patterns may be produced, because the tiling algorithm produces patterns with a nucleotide level resolution. In highly heterogeneous samples, nearly every sequence read may produce a unique tile pattern. Therefore, additional computational analysis is necessary to group similar patterns and classify them into distinct structural variant categories.

**Table 1.**
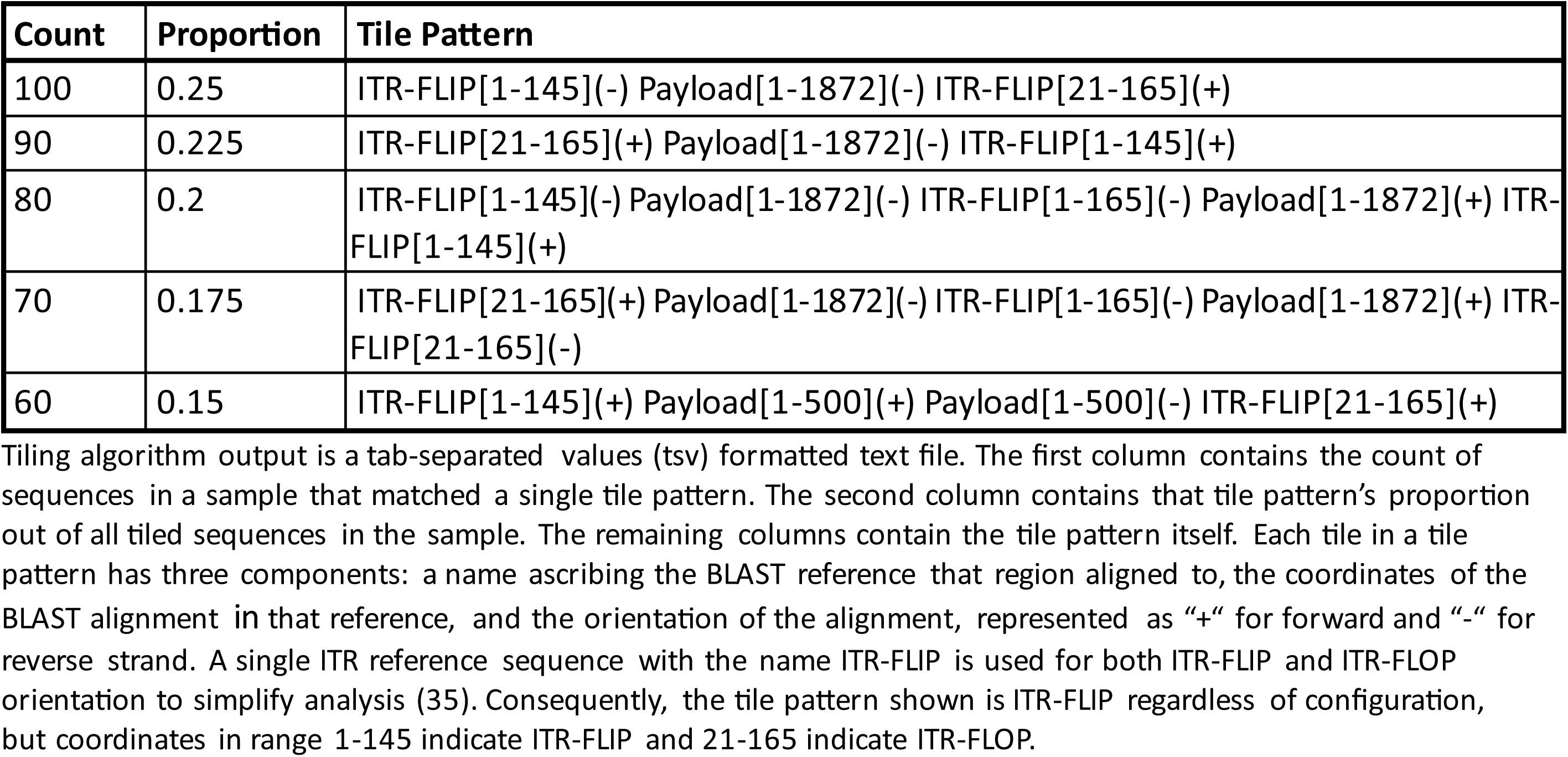
Example tiling algorithm output data.

| Count | Proportion | Tile Pattern |
| --- | --- | --- |
| 100 | 0.25 | ITR-FLIP[1-145](-) Payload[1-1872](-) ITR-FLIP[21-165](+) |
| 90 | 0.225 | ITR-FLIP[21-165](+) Payload[1-1872](-) ITR-FLIP[1-145](+) |
| 80 | 0.2 | ITR-FLIP[1-145](-) Payload[1-1872](-) ITR-FLIP[1-165](-) Payload[1-1872](+) ITR-FLIP[1-145](+) |
| 70 | 0.175 | ITR-FLIP[21-165](+) Payload[1-1872](-) ITR-FLIP[1-165](-) Payload[1-1872](+) ITR-FLIP[21-165](-) |
| 60 | 0.15 | ITR-FLIP[1-145](+) Payload[1-500](+) Payload[1-500](-) ITR-FLIP[21-165](+) |

Tiling algorithm output is a tab-separated values (tsv) formatted text file. The first column contains the count of sequences in a sample that matched a single tile pattern. The second column contains that tile pattern's proportion out of all tiled sequences in the sample. The remaining columns contain the tile pattern itself. Each tile in a tile pattern has three components: a name ascribing the BLAST reference that region aligned to, the coordinates of the BLAST alignment in that reference, and the orientation of the alignment, represented as "+" for forward and "-" for reverse strand. A single ITR reference sequence with the name ITR-FLIP is used for both ITR-FLIP and ITR-FLOP orientation to simplify analysis (35). Consequently, the tile pattern shown is ITR-FLIP regardless of configuration, but coordinates in range 1-145 indicate ITR-FLIP and 21-165 indicate ITR-FLOP.

#### 2.4.2 Subparsing

Using the tiling results, we calculated the count of sequences with contaminant DNA in a sequencing run by counting the tile patterns with any tile that isn’t representing an alignment to the AAV reference sequence. We then analyzed sequences consisting exclusively of canonical AAV reference DNA tiles using our subparser program. The subparser operates in three main stages: lexing, parsing, and final verification. During the lexing step, the names of each alignment in a tile pattern are converted into a set of tokens, which are a finite number of strings in a context-free language (CFL). During the parsing step, any number of tokens produced by our subparser’s lexer are reduced into a single terminal token using a context-free grammar (CFG). This terminal token will be the tile pattern’s tentative structural variant classification. During the final-check step, the orientation and coordinate data of the tile pattern is checked to ensure the accuracy of the tentative structural variant classification. The final structural variant classification is then assigned to the tile pattern. Our subparser program is available on GitHub.

The conversion of the tile pattern names into a CFL, a set of tokens that are readable by a CFG-utilizing parser, is done by a program called a lexer. The vector subparser’s lexer module is built off the ply.lex module available on GitHub (36). The subparser’s lexer converts Payload and ITR-FLIP tile names into “P” and “I” tokens, respectively. The lexer treats both flip and flop ITRs, as well as any other ITR sequence variants, equivalently under the “I” token category to ensure consistent representation across all possible orientations. Single elongated ITRs may be represented by more than one adjacent ITR tile from the tiling program. Consequently, adjacent “I” tokens are reduced to a single “I” token to simplify the CFG and better represent the underlying vector genome structure. An “AND” token is added between each “P” and “I” token to allow for parsing of patterns of any length. These “P”, “I”, and “AND” tokens form the structured CFL input that is passed to the subparser’s parser module for tentative structural variant classification.

The reduction of any number of tokens into a single token by following rules in a CFG (Fig. 2) is done by a program called a parser. The vector subparser’s parser module is built off the ply.yacc module available on GitHub (36). The parser does this by using a left-right, look-ahead (LRLA) parsing algorithm. As the parser proceeds from left to right, the tokens are checked against the rules in the right-hand side of the CFG (Fig. 2). If the tokens match a rule, they are reduced into the token on the left-hand side of that rule.

**Fig 2.**
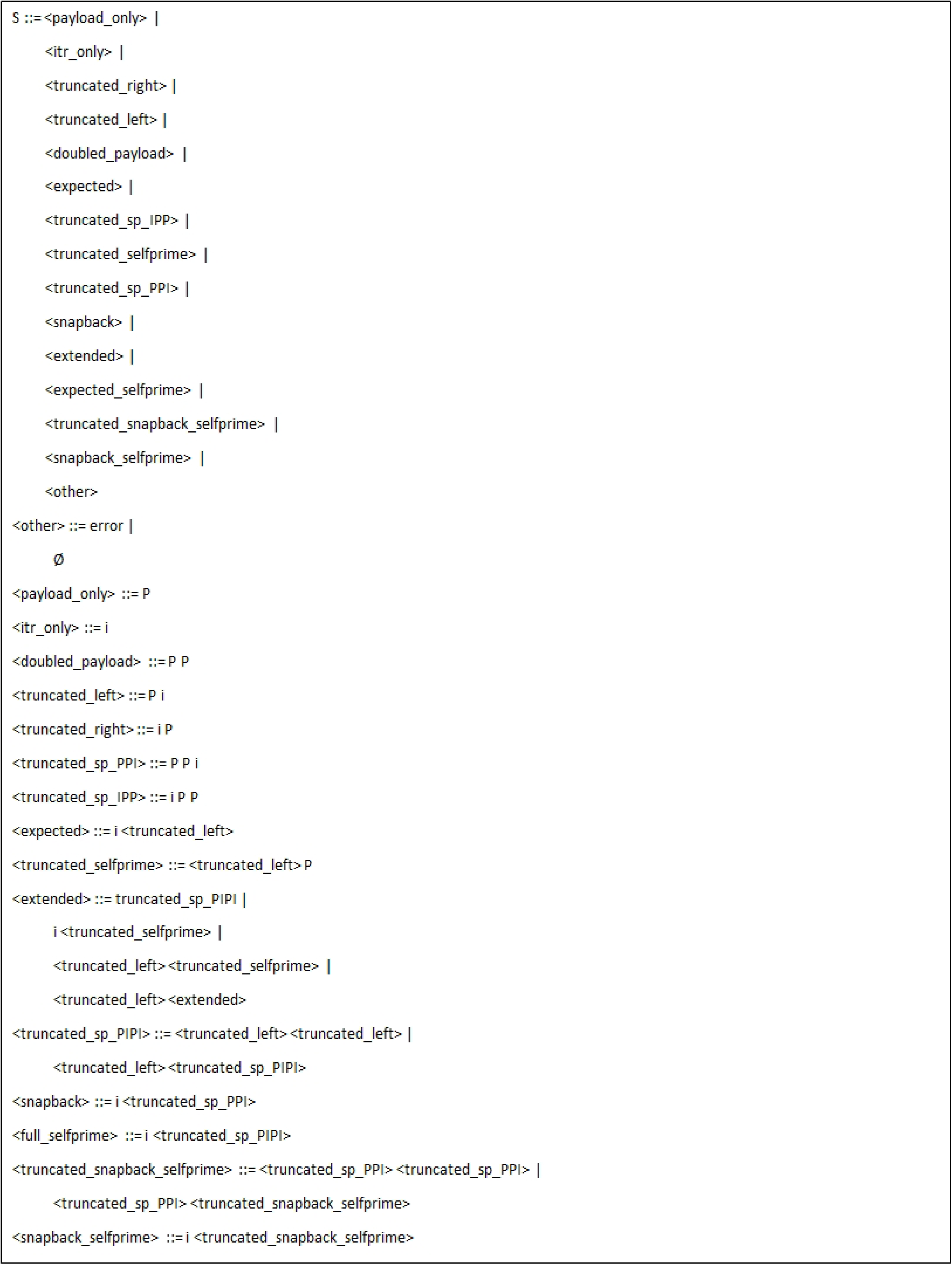
The context-free grammar (CFG) utilized by our subparser program in Backus–Naur form Tokens in this grammar are ”P“, ”I” and “AND”. From the tile pattern, tiles that mapped to the reference AAV payload are converted to the ”P” token, tiles that mapped to the reference ITRs are converted to the” I” token. An ”AND” token is inserted between all adjacent ”I” and ”P” tokens for right-handed precedence. Tokens within inequality signs are terminal tokens and are accepted as the tile pattern structural variant call if they are the only remaining token.

The parser treats the “AND” token as an operator with right-hand precedence, and consequently all tokens will be checked together before any reductions occur. This avoids ambiguous CFG rules by giving longer rules precedence over shorter rules. For example, the rule *<TRUNCATED_SP_PPI> ::= I AND P AND P* will take precedence over the rule *<TRUNCATED_RIGHT> ::= I AND P*, since the former would be checked first. Recursive grammar rules such as *<TRUNCATED_SP_PIPI> ::= <TRUNCATED_LEFT> AND <TRUNCATED_SP_PIPI>* allow for parsing theoretically infinitely long, repeating tile patterns. If the parser is unable to reduce the tokens into a single token, then the parser sets the tile pattern’s tentative structural variant designation to *’other’*.

Since several structural variant calls categorized by our subparser are defined by properties outside of the scope of its parser module’s CFG, including tile orientations and coordinates, a final-check step is done on all tile patterns. For example, an expected structural variant designation necessitates checking the coordinates of the vector genome’s payload sequence to ensure it matches the expected payload sequence. To this end, our subparser checks the tile patterns of sequences given the tentative ‘expected’ structural variant designation to ensure that their payload tile has coordinates matching the expected payload sequence size. If any does not, its structural variant designation is changed to ‘irregular_payload’. The range which a tile pattern’s payload tile start and end coordinates must be in comparison to the reference for it to be considered matching is plus or minus 6bp by default, but this can be modified by the user on the command line. Another example is the check done for all patterns categorized as ‘snapback’ which validates that snapback payloads originate from opposite strands. A complete list of checks that are made after parsing and the full details of the command line options for our subparser program are both included in the supplemental file (S1 File).

### 2.5. *In Silico* Data Generation

To determine the accuracy of the structural variant calling pipeline, we wrote a sequence generation program that takes as input an AAV genome sequence, a configuration file, and three sets of frequency distributions and outputs a file of simulated AAV sequence reads. For the AAV genome sequence, we designed an AAV genome using publicly available sequence data. Each row of the configuration file gives instructions for one run of the sequence generation program, resulting in one simulated sequence file for each row. These rows denote how to organize payload and ITR sequences in the output file, as to simulate the different structural variants callable by our subparser. For example, one row would give the parameters needed to generate sequences with one ITR then one payload to simulate a truncation product missing its second ITR.

To have these generated sequence files better simulate real long-read sequencing data and to robustly test our structural variant calling pipeline, our sequence generator was designed to introduce up to three different kinds of variation to each sequence generated, depending on applicability. To simulate the error rates of different long read sequencers, we generated four different sequence files with a chance of each generated base written being incorrect at a rate of 0, 0.001, 0.01 and 0.05 for each reduction rule in our subparser’s CFG (37–42). To simulate homopolymer and snapback variations seen in real long read sequencing data (14,43), we used the three input frequency distribution sets calculated from real PacBio SMRT NGS sequencing runs. The first set was a single distribution of the frequency of a mutation for each homopolymer size. The second set of frequency distributions was the frequency of each indel size for each homopolymer size. The last set was a single frequency distribution of different lengths of the two payload sequences in a snapback, normalized to reference payload size. The sequence generation program, the generated sequence files, and the frequency distributions used are available on GitHub. Further information on our *in silico* data generation program is included in the supplemental file (S1 File).

### 2.6. Breakpoint-Associated Structure Predictions in Snapback and Truncation Events

To better understand the structural basis for vector genome heterogeneity, particularly within snapback and truncation structural variants, we developed a custom Python–R-based pipeline that performs breakpoint detection followed by local RNA secondary structure analysis. This pipeline operates on subparser output files, which contain sequence-level tile patterns representing the arrangement of vector components such as ITRs and payloads along with their positions and strand orientations.

We parsed these tile patterns using regular expressions to identify genomic breakpoint positions where the continuity of payload segments is interrupted. For snapback structures, we defined the breakpoints as the 3’ nucleotide position in the 5’ payload tile where two payload segments appear in opposite orientations and are flanked on both ends by ITRs (Fig B in S1 File). For self-priming truncation structures, where a single ITR is flanked by two reverse-complementing partial payloads, breakpoints were defined as positions where the flanking payloads were truncated during incomplete replication (Fig C in S1 File). For snapback and self-priming truncation structures, all breakpoint positions were mapped and reported separately for the plus and minus strands.

To investigate the potential structural context of observed breakpoints, we extracted sequence windows surrounding each breakpoint from the reference payload sequence. A range of window sizes was used to capture secondary structure features. Each sequence fragment was folded using RNAfold (v2.5.1, ViennaRNA package(44) with the parameters -p -d2 --noLP to compute the minimum free energy (MFE) and centroid structures. For higher reliability, we retained only those breakpoint-associated windows in which the MFE and centroid structures were identical, indicating consistency in the predicted folding. In cases where multiple overlapping structures were present for the same breakpoint, the structure window with the most negative MFE was selected as the representative conformation for downstream analysis. The final output of this analysis included the breakpoint position, associated sequence window, and corresponding MFE values. To visualize these results, strand-separated plots were generated using ggplot2 package in R (45), showing both the frequency of breakpoints and their associated structural stability across the genome. This approach enabled a comparative view of folding potential around different structural variant junctions within AAV vector preparations.

To further assess sequence features at breakpoint sites, we applied a chi-square goodness-of-fit test using the chi-square from the *SciPy* statistical package (v1.9.3) implemented in Python (v3.10.12). This analysis compared observed nucleotide frequencies at breakpoints positions to the background base composition of the reference payload sequence. Analyses were performed separately for the plus and minus strands.

## 3. Results

### 3.1. *In Silico* Testing

To validate the accuracy of our structural variant calling pipeline, we used our own sequence generator program to produce 68 files of 10,000 AAV genome structural variants sequences and ran them through our structural variant calling pipeline. The files consisted of four replicates of 17 structural variant sequence files, each with a different level of error applied to its sequences. We ran these files through our structural variant calling pipeline and complied the subparser results into a single data table with one row for each file comparing the percentage of sequences in each file that matched its expected structural variant category (S2 Table). We then calculated the average and standard deviation of sequences that matched their expected structural variant category across all 68 files. We then did this same calculation for different subsets of files to investigate pipeline accuracy when different variations were applied to the generated sequences (Table 2).

**Table 2.** *In silico* subparsing summary statistics.

| File Subset | Average Match Percentage | Standard Deviation |
| --- | --- | --- |
| All Files | 99.37% | 0.56% |
| Error Rate 0 | 99.51% | 0.39% |
| Error Rate 0.001 | 99.51% | 0.41% |
| Error Rate 0.01 | 99.45% | 0.44% |
| Error Rate 0.05 | 99.01% | 0.80% |
| Snapbacks | 99.10% | 0.58% |
| Self-Primes | 98.76% | 0.77% |

For all files, the average percentage of sequences in each file that matched their expected structural variant was 99.40% with a standard deviation of 0.56%. For the sequence generation error rates of 0, 0.001, 0.01 and 0.05, the average match percentage and standard deviation across all structural variants were 99.51% and 0.39%, 99.51% and 0.41%, 99.45% and 0.44%, and 99.01% and 0.80% respectively. The average match percentage across all snapback structural variant sequence files which had randomized snapback breakpoint sites was 99.10% with a standard deviation of 0.58%. The average match percentage across all self-priming sequence files was 98.76% with a standard deviation of 0.77%.96.79% for the self-priming expected sequence file with an error rate of 5%. The average match percentage across all self-priming sequence files was 98.76% with a standard deviation of 0.77%.

The minimum match percentage was 96.79% for the self-priming expected sequence file with an error rate of 5%, which is expected due to how we generated homopolymer indels. This was expected because 1) whether or not an indel was generated on a homopolymer was randomized based on frequency distributions, 2) the length generated indels was randomized based on frequency distributions, 3) longer sequences have more homopolymers, and 4) uncommon large indels seen in our frequency distributions prevent BLAST alignment in the tiling step. Consequently, longer sequences like self-priming sequences are more likely to not match their expected category when generated by the sequence generation program.

### 3.2. *IN VITRO* TESTING

To evaluate the consistency and improved resolution of our updated pipeline, we compared the subparser’s output to AUC results for two different AAV samples (Table 3).

**Table 3.** Data comparison between our subparser program utilizing a CFG and orthogonal AUC data for scAAV and ssAAV samples.

| Vector Sample Type | Classification | Percentage | AUC Peak | Percentage |
| --- | --- | --- | --- | --- |
| scGFP | expected | 94.52% | Full | 94.81% |
| scGFP | snapback | 1.97% | Other | 5.19% |
| scGFP | truncated | 1.74% |  |  |
| scGFP | truncated snapback | 0.01% |  |  |
| scGFP | other canonical | 0.56% |  |  |
| scGFP | non-VG | 0.81% |  |  |
| scGFP | unclassified | 0.39% |  |  |
| scGFP | Total | 100.00% | Total | 100.00% |
| ssGFP | expected | 88.59% | Full | 85.79% |
| ssGFP | truncated | 4.38% | Other | 14.21% |
| ssGFP | snapback | 2.50% |  |  |
| ssGFP | truncated snapback | 0.00% |  |  |
| ssGFP | other canonical | 0.79% |  |  |
| ssGFP | non-VG | 3.14% |  |  |
| ssGFP | unclassified | 0.59% |  |  |
| ssGFP | Total | 100.00% | Total | 100.00% |

AUC data is renormalized to exclude empty capsids. non-VG was calculated as a sum of sequences which had any tile in their tile pattern with an alignment to a sequence not within the reference AAV genome sequence. Unclassified sequences were calculated as the difference between a fasta file's sequence count and the sum of the first column of its tiling algorithm output.

For both vector types, the overwhelming majority of genomes were classified as expected full-length species, in close agreement with the full capsid proportions measured by AUC (94.5% vs. 94.8% for scGFP and 88.6% vs. 85.8% for ssGFP). Only small fractions of reads were assigned to noncanonical forms such as snapback (≈2% in both samples) and truncated genomes (1.7% in scGFP and 4.4% in ssGFP), with “truncated snapback” and “other canonical” species each below 1%.

Reads classified as non-VG and unclassified sequences were also very low (<1% for scGFP and ≈3–0.6% for ssGFP). Together, these data show that the subparser’s structural classification produced results highly consistent with the independent AUC measurements, confirming that the pipeline can accurately resolve the major and minor genome populations in rAAV samples.

### 3.3. Breakpoint-Associated Structure Predictions

Snapback genomes identified in both scGFP and ssGFP preparations localized to discrete breakpoint sites along the genome rather than being randomly distributed (Fig. 3A and 3B). For scGFP, snapback junctions were highly concentrated on the minus strand, particularly downstream of the promoter and within the GFP payload region, with several positions showing strong enrichment. In contrast, the plus strand contained relatively few, lower-frequency breakpoints (Fig. 3A). For ssGFP, the overall number of breakpoints was lower than in scGFP, but the junctions still occurred at defined sites rather than being broadly dispersed (Fig. 3B).

**Fig 3.**
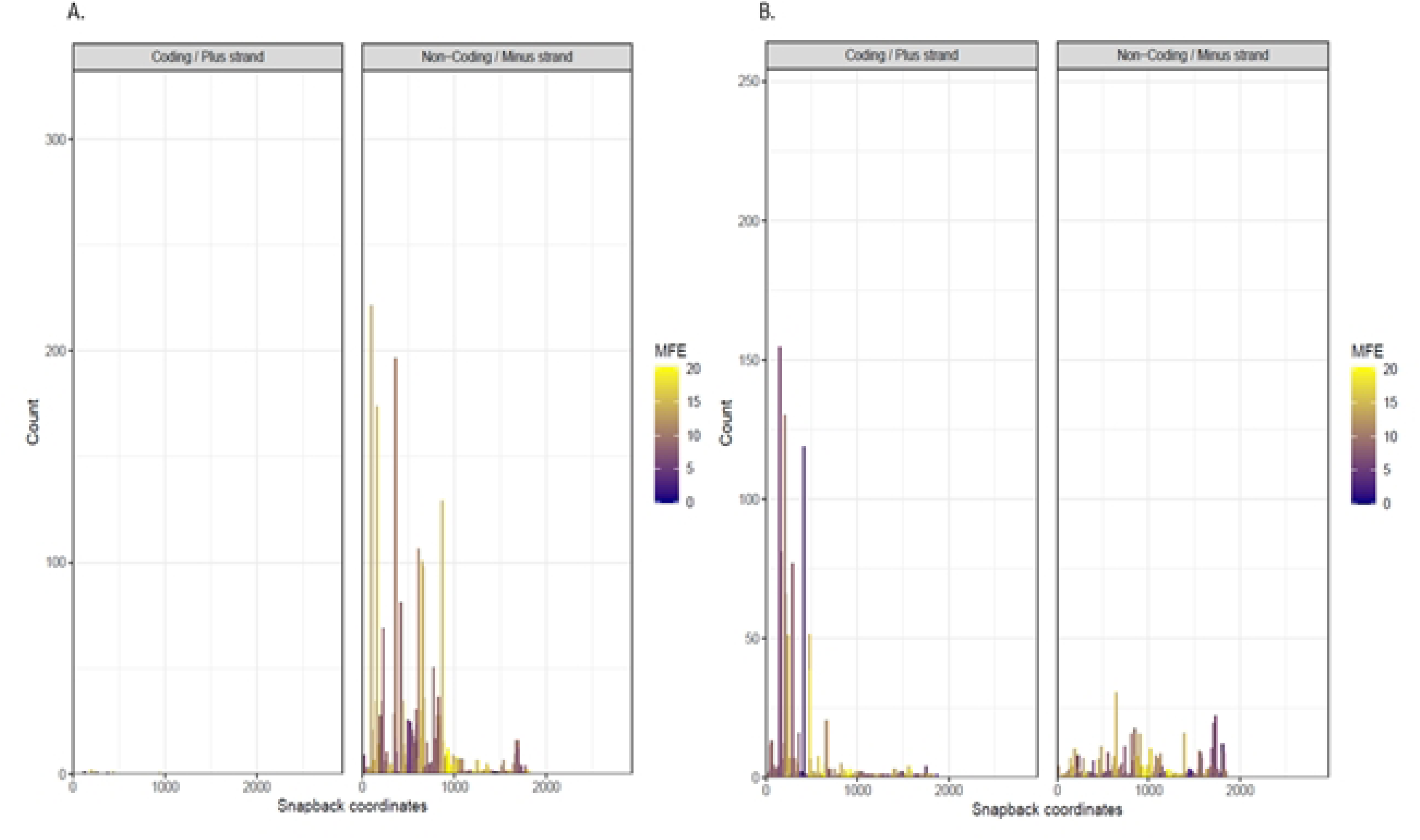
Distribution of Snapback Breakpoints with Local Structural Stability Estimates. A) scGFP B) ssGFP. Bar plots display the count of breakpoints across the AAV payload sequence for plus (coding) and minus (non-coding) strands. The x-axis shows breakpoint positions, and the y-axis indicates the count of reads at each position. Each bar represents a single nucleotide position. Bars are colored according to the absolute value of minimum free energy (MFE) of the predicted RNA secondary structure around each breakpoint, with yellow indicating more stable structures.

To examine potential sequences and structural drivers of these events, we assessed the local folding stability around each breakpoint. Some high-frequency breakpoints coincided with regions predicted to form stable secondary structures (yellow bars in Fig. 3), while others occurred in areas with little or no predicted stability, indicating that secondary structure alone cannot fully account for snapback formation. Despite the presence of significant predicted structure in some snapback nucleotides, several high-frequency breakpoints were also observed in regions with little or no predicted local folding potential, suggesting additional sequence-driven mechanisms.

In addition to snapbacks, we examined truncation-associated self-priming events in both scGFP and ssGFP (Fig D in S1 File). For scGFP, truncation junctions were observed on both strands, with a higher frequency on the minus strand. Several sites clustered at defined positions, whereas others were more broadly distributed across the genome. Some of these breakpoints coincided with stable predicted secondary structures, while many occurred in regions without strong folding potential, again suggesting that both sequence- and structure-driven mechanisms contribute to truncation formation. However, a subset of low-frequency junctions, particularly those with an even genomic distribution, may reflect random DNA shearing introduced during SMRTbell library preparation rather than genuine replication-derived events.

For ssGFP, truncation-associated breakpoints were fewer in number and appeared more dispersed than in scGFP. The relatively low counts and broader distribution suggest that truncation events in ssGFP are less strongly dictated by specific sequence or structure motifs. All identified breakpoints in the scGFP and ssGFP datasets were summarized with their local minimum free energy (MFE), count, and proportional abundance in separate data table files for the snapback and the truncation-associated events (S3-S6 Table).

To integrate these findings, we tested whether sequence composition itself biases breakpoint formation. Nucleotide composition at junctions deviated significantly from background expectations. For snapback genomes, scGFP minus-strand breakpoints were enriched for T (χ² test, p = 8.5×10⁻⁸⁸), whereas the plus strand showed no significant enrichment (p > 0.05). In contrast, ssGFP plus-strand breakpoints were enriched for G and T (χ² test, p = 1.6×10⁻⁷⁰), while the minus strand showed a weaker deviation (p = 3.9×10⁻⁴). In truncation-associated self-priming events, scGFP showed strand-specific bias, with C enrichment on the plus strand (χ² test, p = 7×10⁻⁶) and no significant enrichment on the minus strand (p = 0.14). By contrast, ssGFP truncations showed no strong enrichment on either strand (p > 0.05).

## 4. Discussion

Adeno-associated viral vector (AAV) genome heterogeneity remains a critical concern in gene therapy vector design and quality control. While previous studies have explored the diversity of packaged genomes through various methods, including short-read NGS, they have been unable to reveal the complete structure of entire packaged vector genomes within an AAV sample (46).

To evaluate the robustness and accuracy of our pipeline, we conducted *in silico* benchmarking experiments. The subparser results on the *in silico* data shows that the pipeline was robust in its classifications in the presence of simulated sequencing error rates, homopolymer indel rates, and random snapback breakpoints. Even though classification accuracy declined slightly at the highest simulated error rates and with self-priming sequences, it still correctly classified more than 96% of reads. In the presence of high simulated error rates and random snapback breakpoints, the average match percentage remained over 97%. This consistency suggests that the grammar-based approach can handle the types of artifacts typically seen in long-read sequencing without misclassifying genuine structural variants. In practice, this level of robustness strengthens confidence that differences observed in real AAV samples reflect true biological heterogeneity rather than sequencing noise, making the pipeline suitable for routine quality assessments and comparative studies.

The reduction in accurate calls by our subparser on the *in silico* self-priming sequences was likely due to how our sequence generator program was designed to simulate homopolymer-associated indel errors observed in real sequencing results. This is supported by the lowest match percentage being in the expected self-priming sequence files, which also required payloads that mostly matched the size of the reference AAV genome’s payload sequence. Since these payloads were more likely to contain indels, it was more likely that these sequences failed this check. Even with this expected payload check, with the possibility of large indels being generated, and with a simulated error rate of 5%, the subparser still accurately categorized 96.79% of generated sequences. These results support the subparser’s ability to correctly classify sequences in the presence of indel sequencing errors seen in NGS data.

For both the scGFP sample data and the ssGFP sample data, the subparser had an expected sequence percent comparable to the AUC assay’s full mass peak. Small differences between expected peak size and AUC full mass peak could be a result of non-canonical vector genome structural variants with sizes similar to the expected vector genome size. Despite these differences, the correlation between the AUC results and our NGS pipeline results demonstrates that the pipeline accurately classifies long-read sequencing data. While AUC provides robust, orthogonal estimates of populations ( e.g. full, partial, and empty capsids), it does not reveal sequence-level information. In particular, AUC cannot localize breakpoints, report strand orientation, identify sequence context or secondary-structure features, or distinguish among different canonical structural variants that share similar mass. Our read-level approach fills these gaps by linking each structural call to exact genomic coordinates and sequence features.

Beyond classification, the output of the subparser was analyzed further to investigate biological insights into the structural variant formation process. Our breakpoint analysis provides important insights into the mechanisms underlying snapback and truncation variant formation in AAV genomes. In both scGFP and ssGFP samples, snapback breakpoints were not randomly distributed but instead clustered at discrete genomic sites. Several of these positions overlapped with regions predicted to form stable local secondary structures (low MFE), suggesting that structural folding may promote or stabilize snapback formation. However, other high-frequency breakpoints showed only weak folding potential, indicating that additional processes such as replication stress or non-homologous end joining (NHEJ) may also contribute. Interestingly, similar nonrandom clustering of snapback junctions and nucleotide enrichment patterns was previously described , where G/C nucleotide enrichment at major breakpoints was identified across multiple AAV vector genomes. Our results extend these findings by showing that, in scGFP and ssGFP samples, the specific enrichment pattern varies, suggesting that the preferred nucleotide composition at breakpoints may depend on vector-specific sequence context.

In contrast, truncation events displayed a more uniform and dispersed distribution. The truncation junctions were fewer and appeared more scattered across the genome, consistent with a more stochastic origin. These patterns suggest that truncation products may arise largely from random fragmentation or replication-related instability. Notably, by mapping breakpoints and examining their nearby genomic context, our pipeline can highlight features such as promoters or repetitive elements that may help explain structural variant formation. Furthermore, the pipeline can also be used for the estimation of the sizes of snapback and truncated genomes directly from read-level data, offering insight into the prevalence of shorter genome fragments. In wild-type AAV, such shorter species have been reported to play regulatory roles(43), whereas in recombinant AAV gene therapy vectors, they are best regarded as heterogeneous by products or contaminants. Overall, our findings suggest that secondary structure is one contributing factor but not the only one in the formation of structural variants in AAV vectors. High-resolution structural variant classification, strand-aware breakpoint analysis provides a framework for understanding the diversity of vector genome forms and offers a path to optimizing vector design, helper components, and production conditions.

## 5. Conclusions

Developing long-read sequencing technologies has given researchers the ability to fully sequence vector genomes. Although numerous structural variants have been reported in the literature, there remains a limited number of bioinformatics workflows specifically designed to comprehensively analyze rAAV genome heterogeneity. Using *in silico* and real world sequencing data and by comparing our pipeline results to AUC results, we have demonstrated that our structural variant calling pipeline can accomplish comprehensive characterization of rAAV vectors robustly and accurately. Additionally, the tile patterns that match each structural variant can be written into separate files so that downstream analysis can be restricted to only sequences with a desired morphology.

To complement structural variant calling and to demonstrate analysis of sequences filtered by our subparser program, we also developed a structural analysis module that examines strand-specific breakpoints and local RNA folding potential. This adds mechanistic insight into how sequence structure may influence genome heterogeneity, especially in snapback and truncation variants.

Overall, this pipeline advances the resolution at which vector genome complexity can be studied and understood. Its application across ssAAV and scAAV samples and its correlation with AUC results demonstrates its practical utility for both research and vector quality control.

## Supporting Information

S1 File. Supplemental Figures, Descriptions, and Tables.

S2 Table. *In Silico* Subparsing Results.

S3 Table. Snapback scGFP results.

S4 Table. Snapback scGFP results.

S5 Table. Truncation self-priming scGFP results.

S6 Table. Truncation self-priming ssGFP results.

## Data Availability

The full codebase is available on GitHub at https://github.com/drouleau0/rAAV_Structural_Variant_Classifier to support reproducibility and community use.

## Acknowledgment

We thank James McGivney for his support in establishing AAV sequencing capabilities. We are also grateful to Kiran Adhikari and Celia Slater for their assistance in reviewing the manuscript, and to Brenda Burnham for performing the AUC analyses. We further acknowledge the team at Pacific Biosciences for their expert technical assistance and for ensuring reliable performance of the Sequel II sequencing platform.

## References

1. Fong S, Yates B, Sihn C-R, Mattis AN, Mitchell N, Liu S, et al. Interindividual variability in transgene mRNA and protein production following adeno-associated virus gene therapy for hemophilia A. Nat Med. 2022 Apr;28(4):789–97.

2. Russell S, Bennett J, Wellman JA, Chung DC, Yu Z-F, Tillman A, et al. Efficacy and safety of voretigene neparvovec (AAV2-hRPE65v2) in patients with RPE65 -mediated inherited retinal dystrophy: a randomised, controlled, open-label, phase 3 trial. Lancet. 2017 Aug;390(10097):849–60.

3. Kuzmin DA, Shutova MV, Johnston NR, Smith OP, Fedorin VV, Kukushkin YS, et al. The clinical landscape for AAV gene therapies. Nat Rev Drug Discov. 2021 Mar 25;20(3):173 –4.

4. Duan D. Lethal immunotoxicity in high-dose systemic AAV therapy. Mol Ther. 2023 Nov 1;31(11):3123–6.

5. Salabarria SM, Corti M, Coleman KE, Wichman MB, Berthy JA, D’Souza P, et al. Thrombotic microangiopathy following systemic AAV administration is dependent on anti-capsid antibodies. J Clin Invest [Internet]. 2024 Jan 2;134(1). Available from: 10.1172/JCI173510

6. Silver E, Argiro A, Hong K, Adler E. Gene therapy vector-related myocarditis. Int J Cardiol. 2024 Mar 1;398(131617):131617.

7. de Jong YP, Herzog RW. Liver gene therapy and hepatocellular carcinoma: A complex web. Mol Ther. 2021 Apr 7;29(4):1353–4.

8. Ertl HCJ. Immunogenicity and toxicity of AAV gene therapy. Front Immunol. 2022 Aug 12;13:975803.

9. FDA Recommends Removal of Voluntary Hold for Elevidys for ambulatory patients [Internet]. U.S. Food and Drug Administration. FDA; 2025 [cited 2025 Sept 4]. Available from: https://www.fda.gov/news-events/press-announcements/fda-recommends-removal-voluntary-hold-elevidys-ambulatory-patients

10. Gross D-A, Tedesco N, Leborgne C, Ronzitti G. Overcoming the challenges imposed by humoral immunity to AAV vectors to achieve safe and efficient gene transfer in seropositive patients. Front Immunol. 2022 Apr 7;13:857276.

11. Rosas LE, Grieves JL, Zaraspe K, La Perle KM, Fu H, McCarty DM. Patterns of scAAV vector insertion associated with oncogenic events in a mouse model for genotoxicity. Mol Ther. 2012 Nov;20(11):2098 – 110.

12. Xie J, Mao Q, Tai PWL, He R, Ai J, Su Q, et al. Short DNA hairpins compromise recombinant adeno-associated virus genome homogeneity. Mol Ther. 2017 June 7;25(6):1363 –74.

13. Namkung S, Tran NT, Manokaran S, He R, Su Q, Xie J, et al. Direct ITR-to-ITR nanopore sequencing of AAV vector genomes. Hum Gene Ther. 2022 Nov;33(21–22):1187–96.

14. Zhang J, Guo P, Yu X, Frabutt DA, Lam AK, Mulcrone PL, et al. Subgenomic particles in rAAV vectors result from DNA lesion/break and non-homologous end joining of vector genomes. Mol Ther Nucleic Acids. 2022 Sept 13;29:852–61.

15. McColl-Carboni A, Dollive S, Laughlin S, Lushi R, MacArthur M, Zhou S, et al. Analytical characterization of full, intermediate, and empty AAV capsids. Gene Ther. 2024 May 19;31(5 –6):285–94.

16. Barnes LF, Draper BE, Kurian J, Chen Y-T, Shapkina T, Powers TW, et al. Analysis of AAV-extracted DNA by charge detection mass spectrometry reveals genome truncations. Anal Chem. 2023 Mar 7;95(9):4310–6.

17. Tran NT, Lecomte E, Saleun S, Namkung S, Robin C, Weber K, et al. Human and insect cell-produced recombinant adeno-associated viruses show differences in genome heterogeneity. Hum Gene Ther. 2022 Apr;33(7–8):371–88.

18. Zhang J, Yu X, Guo P, Firrman J, Pouchnik D, Diao Y, et al. Satellite subgenomic particles are key regulators of adeno-associated virus life cycle. Viruses. 2021 June 21;13(6):1185.

19. Tran NT, Heiner C, Weber K, Weiand M, Wilmot D, Xie J, et al. AAV-genome population sequencing of vectors packaging CRISPR components reveals design-influenced heterogeneity. Mol Ther Methods Clin Dev. 2020 Sept 11;18:639–51.

20. Barnes LF, Draper BE, Chen Y-T, Powers TW, Jarrold MF. Quantitative analysis of genome packaging in recombinant AAV vectors by charge detection mass spectrometry. Mol Ther Methods Clin Dev. 2021 Dec 10;23:87–97.

21. Zhang J, Chrzanowski M, Frabutt DA, Lam AK, Mulcrone PL, Li L, et al. Cryptic resolution sites in the vector plasmid lead to the heterogeneities in the rAAV vectors. J Med Virol. 2023 Jan;95(1):e28433.

22. Gimpel AL, Katsikis G, Sha S, Maloney AJ, Hong MS, Nguyen TNT, et al. Analytical methods for process and product characterization of recombinant adeno-associated virus-based gene therapies. Mol Ther Methods Clin Dev. 2021 Mar 12;20:740–54.

23. Lecomte E, Tournaire B, Cogné B, Dupont J-B, Lindenbaum P, Martin-Fontaine M, et al. Advanced characterization of DNA molecules in rAAV vector preparations by single-stranded virus next-generation sequencing. Mol Ther Nucleic Acids. 2015 Oct 27;4(e260):e260.

24. Guerin K, Rego M, Bourges D, Ersing I, Haery L, Harten DeMaio K, et al. A novel next-generation sequencing and analysis platform to assess the identity of recombinant adeno-associated viral preparations from viral DNA extracts. Hum Gene Ther. 2020 June;31(11–12):664–78.

25. Center for Biologics Evaluation, Research. Recommendations for Microbial Vectors Used for Gene Therapy [Internet]. U.S. Food and Drug Administration. FDA; 2019 [cited 2025 Mar 13]. Available from: https://www.fda.gov/regulatory-information/search-fda-guidance-documents/recommendations-microbial-vectors-used-gene-therapy

26. 26. Talevich E, Tseng E, Diallo A, Sellami N, Elliott A, Cantarel BL, et al. Standardized nomenclature and reporting for PacBio HiFi sequencing and analysis of rAAV gene therapy vectors [Internet]. bioRxiv. 2024. p. 2024.05.07.592296. Available from: http://biorxiv.org/content/early/2024/05/10/2024.05.07.592296.abstract

27. Bruccoleri RE, Rouleau D, Slater C, Lata D, Phillion C, Adjei S, et al. The Tiling Algorithm – A general method for structural characterization of accurate long DNA sequence reads: application to AAV genome sequences [Internet]. bioRxiv. 2025. Available from: 10.1101/2025.07.25.666743

28. Altschul SF, Gish W, Miller W, Myers EW, Lipman DJ. Basic local alignment search tool. J Mol Biol. 1990 Oct 5;215(3):403–10.

29. Ferrari FK, Samulski T, Shenk T, Samulski RJ. Second-strand synthesis is a rate-limiting step for efficient transduction by recombinant adeno-associated virus vectors. J Virol. 1996 May;70(5):3227–34.

30. [cited 2025 Sept 8]. Available from: https://www.pacb.com/wp-content/uploads/Procedure-checklist-Preparing-multiplexed-AAV-SMRTbell-libraries-using-SMRTbell-prep-kit-3.0.pdf

31. Troxell B, Jaslow SL, Tsai I-W, Sullivan C, Draper BE, Jarrold MF, et al. Partial genome content within rAAVs impacts performance in a cell assay-dependent manner. Mol Ther Methods Clin Dev. 2023 Sept 14;30:288–302.

32. Zhang J, Yu X, Chrzanowski M, Tian J, Pouchnik D, Guo P, et al. Thorough molecular configuration analysis of noncanonical AAV genomes in AAV vector preparations. Mol Ther Methods Clin Dev. 2024 Mar 14;32(1):101215.

33. Tam Tran N, WL Tai P. Profiling AAV vector heterogeneity & contaminants using next-generation sequencing methods. Cell Gene Ther Insights. 2024 Jan 8;09(11):1565 –83.

34. Werle AK, Powers TW, Zobel JF, Wappelhorst CN, Jarrold MF, Lyktey NA, et al. Comparison of analytical techniques to quantitate the capsid content of adeno-associated viral vectors. Mol Ther Methods Clin Dev. 2021 Dec 10;23:254–62.

35. Wilmott P, Lisowski L, Alexander IE, Logan GJ. A user’s guide to the inverted terminal repeats of adeno-associated virus. Hum Gene Ther Methods. 2019 Dec;30(6):206–13.

36. Beazley D. PLY Python Lex-Yacc [Internet]. 2018 [cited 2024 Mar 8]. Available from: https://github.com/dabeaz/ply.

37. Wenger AM, Peluso P, Rowell WJ, Chang P-C, Hall RJ, Concepcion GT, et al. Accurate circular consensus long-read sequencing improves variant detection and assembly of a human genome. Nat Biotechnol. 2019 Oct;37(10):1155–62.

38. Weirather JL, de Cesare M, Wang Y, Piazza P, Sebastiano V, Wang X-J, et al. Comprehensive comparison of Pacific Biosciences and Oxford Nanopore Technologies and their applications to transcriptome analysis. F1000Res. 2017 Feb 3;6(100):100.

39. Wang Y, Zhao Y, Bollas A, Wang Y, Au KF. Nanopore sequencing technology, bioinformatics and applications. Nat Biotechnol. 2021 Nov;39(11):1348–65.

40. Delahaye C, Nicolas J. Sequencing DNA with nanopores: Troubles and biases. PLoS One. 2021 Oct 1;16(10):e0257521.

41. Goodwin S, Gurtowski J, Ethe-Sayers S, Deshpande P, Schatz MC, McCombie WR. Oxford Nanopore sequencing, hybrid error correction, and de novo assembly of a eukaryotic genome. Genome Res. 2015 Nov 1;25(11):1750–6.

42. Rang FJ, Kloosterman WP, de Ridder J. From squiggle to basepair: computational approaches for improving nanopore sequencing read accuracy. Genome Biol. 2018 July 13;19(1):90.

43. Ross MG, Russ C, Costello M, Hollinger A, Lennon NJ, Hegarty R, et al. Characterizing and measuring bias in sequence data. Genome Biol. 2013 May 29;14(5):R51.

44. Lorenz R, Bernhart SH, Höner Zu Siederdissen C, Tafer H, Flamm C, Stadler PF, et al. ViennaRNA Package 2.0. Algorithms Mol Biol. 2011 Nov 24;6(1):26.

45. Wickham H. Ggplot2. 2nd ed. Cham, Switzerland: Springer International Publishing; 2016. 260 p. (Use R!).

46. Understanding heterogeneity in rAAV products and impurities [Internet]. [cited 2025 Sept 4]. Available from: https://www.darkhorseconsultinggroup.com/post/heterogeneity-in-raav-vector-genomes-etc

